# In vivo characterization and quantification of neurofibrillary tau PET radioligand ^18^F-MK-6240 in humans from Alzheimer’s disease dementia to young controls

**DOI:** 10.1101/290064

**Authors:** Tobey J Betthauser, Karly A Cody, Matthew D Zammit, Dhanabalan Murali, Alexander K Converse, Todd E Barnhart, Charles K Stone, Howard A Rowley, Sterling C Johnson, Bradley T Christian

## Abstract

Tau positron emission tomography (PET) imaging has potential for elucidating changes in the deposition of neuropathological tau aggregates that are occurring during the progression of Alzheimer’s disease (AD). This work investigates *in vivo* kinetics, quantification strategies and imaging characteristics of a novel tau PET radioligand ^18^F-MK-6240 in humans.

**Methods:** Fifty-one individuals ranging from cognitively normal young controls to persons with dementia underwent T1-weighted magnetic resonance imaging (MRI), and ^11^C-PiB and ^18^F-MK-6240 PET imaging. PET data were coregistered to the MRI and time-activity curves were extracted from regions of interest to assess ^18^F-MK-6240 kinetics. The pons and inferior cerebellum were investigated as potential reference regions. Reference tissue methods (Logan graphical analysis (LGA) and multilinear reference tissue method (MRTM2)) were investigated for quantification of ^18^F-MK-6240 distribution volume ratios (DVRs) in a subset of nineteen participants. Stability of DVR methods was evaluated using truncated scan durations. Standard uptake value ratio (SUVR) estimates were compared to DVR estimates to determine the optimal timing window for SUVR analysis. Parametric SUVR images were used to identify regions of potential off-target binding and to compare binding patterns with neurofibrillary tau staging established in neuropathology literature.

**Results:** Standard uptake values in the pons and the inferior cerebellum indicated consistent clearance across all 51 subjects. LGA and MRTM2 DVR estimates were similar, with LGA slightly underestimating DVR compared to MRTM2. DVR estimates remained stable when truncating the scan duration to 60 minutes. SUVR determined 70-90 minutes post-injection of ^18^F-MK-6240 indicated linearity near unity when compared to DVR estimates and minimized potential spill-in from uptake outside of the brain. ^18^F-MK-6240 binding patterns in target regions were consistent with neuropathological neurofibrillary tau staging. Off-target binding regions included the ethmoid sinus, clivus, meninges, substantia nigra, but not the basal ganglia or choroid plexus.

**Conclusions:** ^18^F-MK-6240 is a promising PET radioligand for *in vivo* imaging of neurofibrillary tau aggregates in AD with minimal off-target binding in the human brain.

## INTRODUCTION

### AD Pathophysiology

Neurofibrillary tau tangles (NFTs) and beta-amyloid (Aβ) plaques are neuropathological hallmarks of AD that accumulate decades prior to neurodegeneration and symptomatic onset of disease (1–3). Neuropathological staging of NFTs and Aβ plaques suggests these protein aggregates follow hierarchical spatiotemporal patterns during the AD progression that are indicative of disease severity. Specific to tau, neuropathological staining in predetermined slices indicates NFTs are first observed in the transentorhinal cortex (Braak stage I), followed by the hippocampus (stage II), and then spread laterally to the inferior temporal cortex (stage III) and subsequently spread throughout the neocortex in an ordered pattern (stages IV-VI) (4). *In vivo* biomarker studies evaluating Aβ and tau are mostly consistent with postmortem findings and suggest a temporal biomarker cascade that putatively begins with detectable Aβ accumulation, followed by tau aggregation and ultimately neurodegeneration and deficits in cognition (5–7). Recently, cross-sectional analyses of Aβ and tau PET imaging have indicated these biomarkers follow hierarchical patterns that may be useful for *in vivo* disease staging (7,8). Characterization of the temporal sequencing of the biomarker cascade shows promise for predicting future cognitive decline, particularly at the patient level, which could dramatically improve late-life planning and outcomes of clinical prevention trials (9).

### Tau PET Imaging

Starting in 2013 PET ligands for detecting NFTs have undergone rapid development (for review see (10)). Initial tau PET imaging studies have indicated approximate concordance between *in vivo* imaging (11–13) and hierarchical patterns observed in neuropathological staging of NFT’s. These studies also demonstrate relationships between tracer specific binding and various measures of cognition, neurodegeneration and delineation of clinical groups. These early results suggest tau PET imaging will play a critical role in disentangling the complex interactions between Aβ, neurofibrillary tau, neurodegeneration and their impact on cognition and late-stage disease outcomes (14). While these studies highlight the promise for tau imaging, they primarily focus on comparing clinically impaired individuals to cognitively healthy controls and do not address the potential of these ligands for identifying tau in early-stage disease where disease intervention is likely to be more effective. Additionally, some tau PET ligands have suffered from lack of specificity for tau (THK series) (15,16), nonpolar radiometabolites (PBB3) (17), and off-target binding that interferes with regions of interest for monitoring early-stage tau deposition such as the hippocampus (AV-1451 and THK series) (12). Recently, ^18^F-MK-6240 has shown high *in vitro* affinity for NFTs and no *in vivo* off-target binding in the basal ganglia in non-human primates (18). These preclinical results indicate ^18^F-MK-6240 has potential for selective imaging of AD tau aggregates in humans, and may be more sensitive for detecting tau in regions associated with early NFT deposition (i.e. early Braak regions). The central aims of this work are to 1) evaluate the *in vivo* pharmacokinetics, 2) investigate dynamic and static reference tissue methods for quantification of specific binding, 3) characterize the *in vivo* spatial distribution of binding related to tau, and 4) identify regions of potential off-target binding of ^18^FMK-6240 in humans ranging from cognitively unimpaired young adults to clinically diagnosed probable AD.

## MATERIALS AND METHODS

### Participants and Recruitment

Participants (n=51) were recruited from the University of Wisconsin-Madison Alzheimer’s Disease Research Center and its affiliated clinics, or the Wisconsin Registry for Alzheimer’s Prevention (19), a longitudinal study following late-middle-aged individuals enriched for AD risk. AD dementia individuals were determined based on clinical diagnosis of probable AD (diagnosis was not informed by AD biomarkers). All other participants were grouped as young controls (27-45 years), older controls, cognitive decliners, and mild cognitive impairment (MCI). The latter three diagnoses were based on longitudinal neuropsychological evaluation and consensus diagnosis (19). Descriptive statistics for the groups are summarized in Table 1. Written informed consent was obtained from all individuals prior to participation. This study was conducted under the University of Wisconsin-Madison Institutional Review Board and the Federal Drug Administration Investigational New Drug mechanism for ^18^F-MK-6240 and ^11^C-PiB PET studies. No adverse events were reported for administration of ^18^F-MK-6240.

**TABLE 1.**
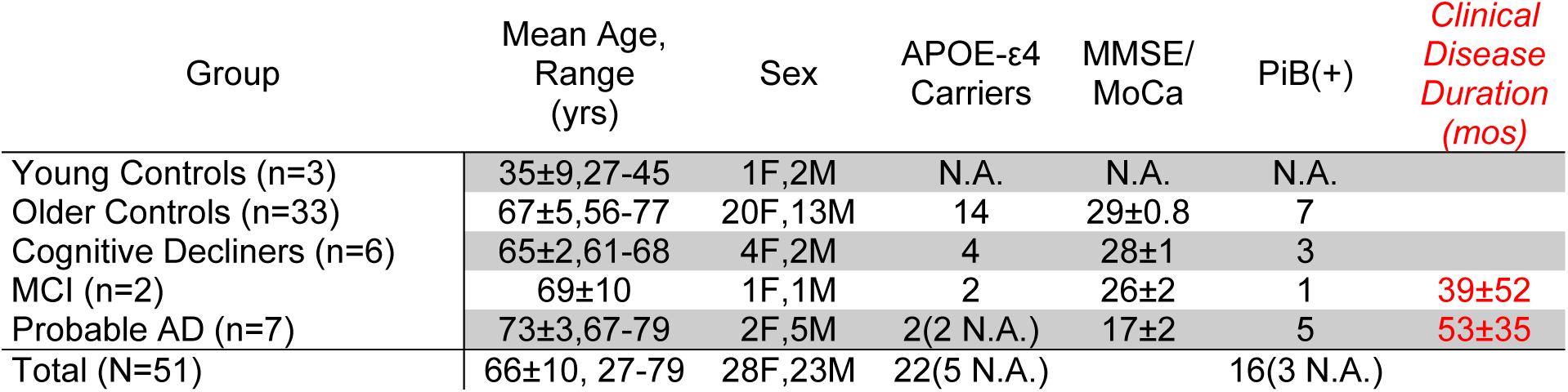
Descriptive statistics for study participants.

### MRI and Anatomical Delineation

All participants underwent a T1-weighted 3D inversion recovery fast spoiled gradient-echo sequence on a 3T MRI scanner (Signa 750, GE Healthcare) with a 32-channel head coil (inversion time, 450 ms; repetition time, 8.1 ms; echo time, 3.2 ms; flip angle, 12°; matrix, 256×256×156; voxel dimensions, 1×1×1 mm; field of view, 256 mm; slice thickness, 1.0 mm). The T1-w image was corrected for magnetic field inhomogeneity (SPM12) and tissue class segmented for white matter (WM), gray matter (GM), and cerebrospinal fluid (CSF). The deformation field obtained from the tissue-class segmentation was used to inverse warp regions of interest (ROIs) from MNI template space to native MRI space.

### Radiochemical Synthesis

^1*1*^*C-PiB* Radiochemical synthesis of ^11^C-PiB was performed as previously described (20) yielding specific activity of 650±161 MBq/nmol (mean±SD, n=48).

*^18^F-MK-6240* ^18^F-MK-6240 was synthesized similar to previously reported methods (18) with modifications to improve ^18^F-MK-6240 radiochemical yield and automated using a Sofie ELIXYS (21) and a computer controlled fraction collection, solid-phase extraction and formulation module previously validated for human use (22). ^18^F-Fluoride was isolated from bulk target ^18^O-water (98% enrichment) after cyclotron irradiation (∼20 μA•h) using an anion exchange column (QMA Accell Plus Light) eluted with 0.8 mL 80/20 acetonitrile/water with 5.6 mg 4,7,13,16,21,24-hex-aoxa-1,10-diazabicyclo[8.8.8]hexacosane (Kryptofix®222) and 1.7 mg potassium carbonate and rinsed with 0.8 mL anhydrous acetonitrile. After three times azeotropic distillation of the ^18^F-KF solution (110 ºC), 1.0 mg MK-6420 precursor dissolved in 650 μL anhydrous dimethyl sulfoxide was added and heated at 140 ºC for 10 minutes. Reaction product was hydrolyzed (3N HCl, 8 minutes at 90 ºC), neutralized with 2.85 mL sodium hydroxide and underwent solid phase extraction (diluted with 2 mL deionized water, tC18 Sep-Pak Plus Light, rinsed with 6 mL deionized water, eluted with 1.15 mL ethanol). The eluate from solid phase extraction was diluted in 1.15 mL 10 mM sodium acetate and was purified via semi-preparative high performance liquid chromatography (Gemini 5 μm C6-phenyl 110 Å 250x10 mm, 45/55 ethanol/10 mM sodium acetate, 3-4 mL/min). The ^18^F-MK-6240 fraction was collected in a bottle containing 35 mL sterile water for injection, USP and underwent solid phase extraction (tC18 Sep-Pak Plus Short, rinsed with 15 mL sterile water for injection, USP, eluted with 1 mL dehydrated ethanol). The eluate was diluted in 9 mL bacteriostatic 0.9% sodium chloride for injection, USP and was 0.22 μm filtered and collected in a vented 10 mL sterile empty vial. A summary of radiochemistry results for ^18^F-MK-6240 syntheses can be found in Table 2.

**TABLE 2.** Summary statistics for ^18^F-MK-6240 radiochemical syntheses and PET radiotracer injection. (NDC=non-decay corrected)

| *Metric (N=51)* | *Mean±SD* |
| --- | --- |
| *Specific Activity (MBq / nmol)* | 852±375 |
| *NDC Yield (%)* | 12.5±4.0 |
| *Synthesis Time (min)* | 101±13 |
| *Injected Activity (MBq)* | 393±7 |
| *Injected Mass (nmol)* | 0.69±0.28 |
| *(range)* | (0.25-1.59) |

### PET Imaging

PET scans were acquired using a Siemens ECAT EXACT HR+ tomograph. *^11^C-PiB* Dynamic ^11^CPiB scans were acquired from 0-70 minutes after a nominal 555 MBq injection for all participants except young controls, who were assumed to be devoid of Aβ pathology based on their age (3). DVRs (LGA, cerebellar GM reference region) were estimated (20) and a global DVR threshold (23) was used to ascertain PiB status (PiB(+) or PiB(−)) for descriptive purposes.

*^18^F-MK-6240* A total of 51 participants underwent ^18^F-MK-6240 PET scans following a nominal 370 MBq injection. A subset of nineteen participants (3 young controls, 6 older controls, 2 cognitive decliners, 1 MCI, 7 probable AD) were scanned dynamically from bolus tracer injection for a total duration of either 90, 105, or 120 minutes. The remaining 32 participants were scanned for 60 minutes following a 60-minute uptake period. ^18^F-MK-6240 PET images were reconstructed using optimized subset expectation maximization (ECAT v7.2.2, 4 iterations, 16 subsets, brain mode on, ramp filter, voxel size 2.57x2.57x2.425 mm, matrix size 128x128x63, corrections applied: segmented attenuation, detector deadtime, scatter, detector normalization and radioisotope decay).

### Data Extraction, Quantification and Analysis of Simplified Methods

The reconstructed ^18^F-MK-6240 PET time series was interframe realigned and coregistered to T1-w MRI (SPM12). HYPR-LR denoising (24) was applied in native PET space to the realigned PET scans with full dynamic data used for DVR analysis (see below). Parametric ^18^F-MK-6240 standard uptake value ratio (SUVR=C(t)/Cref(t)) images were generated using data from 70-90 minutes post injection (Cref, inferior cerebellar GM reference region). ^18^F-MK-6240 time-activity curves (TACs) were extracted from the coregistered PET time series in native T1 space. Pons and off-target ROIs were delineated in MNI space based on an in-depth imaging review of ^18^F-MK-6240 parametric SUVR images in individual cases and SUVR images averaged across control subjects (see below). Additionally, an inferior cerebellum ROI was generated for reference region analysis by combining Automated Anatomical Labeling (AAL, Neurodegenerative Diseases Institute, Université de Bordeaux) ROIs (93,94,101-104) in native T1 space and eroding the mask to limit spill-in from adjacent WM, CSF. The inferior cerebellum (henceforth referred to as “cerebellum”) was used, as opposed to the entire cerebellar GM, due to focal binding observed in the the superior cerebellum and the adjacent tentorium cerebelli, and to avoid contamination from occipital cortex spillover observed in AD participants with high occipital retention. Brain penetrance and evaluation of reference regions (cerebellum and pons) was performed by comparing the standard uptake value (SUV = CPET / ID x mass) across all subjects. DVRs were determined at the ROI level using reference tissue LGA (25) and MRTM2 (26) for all participants with full dynamic acquisitions (n=19). LGA and MRTM2 were chosen as they are amenable to generation of parametric images for voxel-wise analyses. LGA and MRTM2 fitting times (t*) were determined by comparing DVR estimates using stepwise t* values (Bland-Altman plots). Stability of the DVR as a function of scan duration was evaluated within each method by regressing DVR estimates derived from truncated data (i.e. shorter scans) onto DVRs derived from full-length scans. SUVRs derived from stepwise 20-minute windows starting 40 minutes post-injection were regressed onto LGA and MRTM2 DVR to assess the quantitative accuracy of SUVR.

ROIs consisted of the AAL atlas, which was restricted to voxels with GM probabilities greater than thirty percent, and manually segmented ROIs drawn in MNI space in regions with apparent off-target binding. Regions for regression analyses included all AAL ROIs (n=90) except cerebellar regions and also did not include the manually segmented off-target ROIs. This was done to capture the full range of binding observed in this study throughout the entire brain.

### ^18^F-MK-6240 Image Review

Individual parametric ^18^F-MK-6240 SUVR images were reviewed to identify regions of potential off-target binding (not consistent with neuropathology literature) and tau-specific binding (consistent with neuropathology literature) by consensus of a neuroradiologist (Rowley) and experienced neuroimagers blinded to amyloid imaging, cognitive trajectory and clinical group. Additionally, ^18^F-MK-6240 parametric images normalized to MNI space were averaged for PiB(−) controls and PiB(+) AD and MCI cases to aid in the identification of common off-target binding regions and off-target ROI delineation.

## RESULTS

### Reference Region Evaluation

SUV TACs indicated consistent washout across all 51 subjects in the cerebellum and the pons (Fig. 1), and brain penetrance similar to other PET radioligands (peak SUV∼2.5-5). The cerebellum was used as reference region for the remainder of the analyses due to the larger ROI volume (17.1±3.8 cm^3^ vs. 2.2±0.4 cm^3^) and previous validation with other tau PET radioligands(27,28).

**FIGURE 1.**
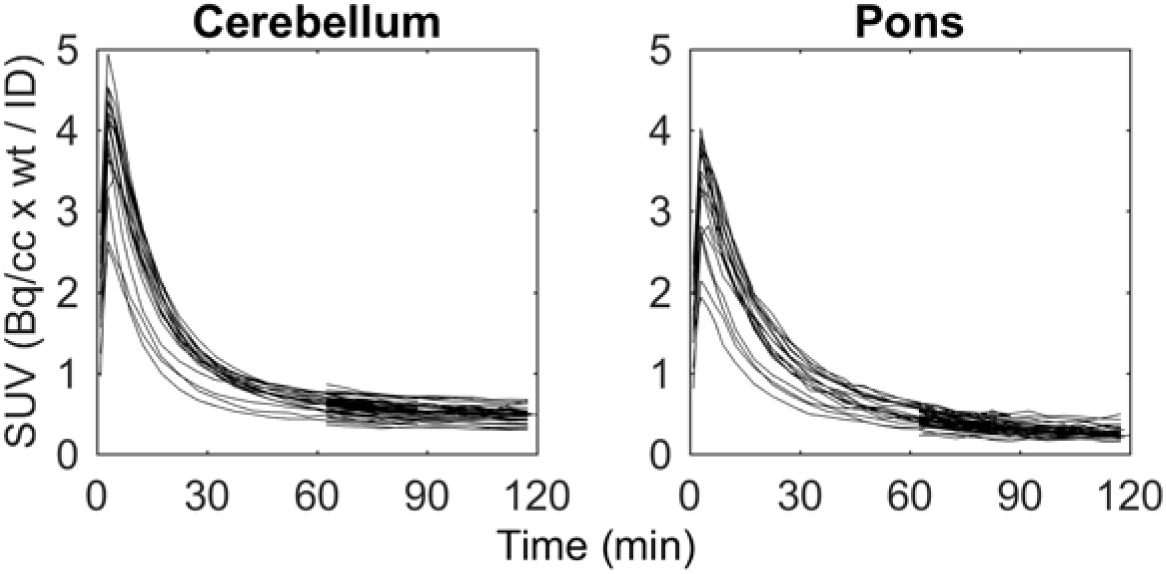
^18^F-MK-6240 standard uptake value time-activity curves of all 51 participants in the cerebellum (left) and the pons (right). No individuals had evidence of specific binding in either region of interest (ROI).

### Pharmacokinetic Evaluation of ^18^F-MK-6240

Target-to-cerebellum TACs (Fig. 2) plateaued around 70 minutes post injection for moderate binding subjects and regions, but were still increasing at 90 minutes in neocortical regions of the highest binding AD subjects (SUVR>3). TACs relative to cerebellum in off-target regions that included bone marrow (ethmoid sinus, clivus, sphenotemporal buttress) were increasing throughout the entire 120-minute scan duration (Fig. 2) and had SUVR values similar to the parahippocampus and the inferior temporal gyrus of PiB(+) AD and MCI individuals around 90 minutes. Similarly, TACs in the meninges relative to cerebellum were increasing throughout the entire 120-minute scan and varied considerably in magnitude across subjects (SUVR 0.5-3.5 at 120 minutes). High meninges binding was more frequent in individuals that did not exhibit specific binding in NFT associated regions.

**FIGURE 2.**
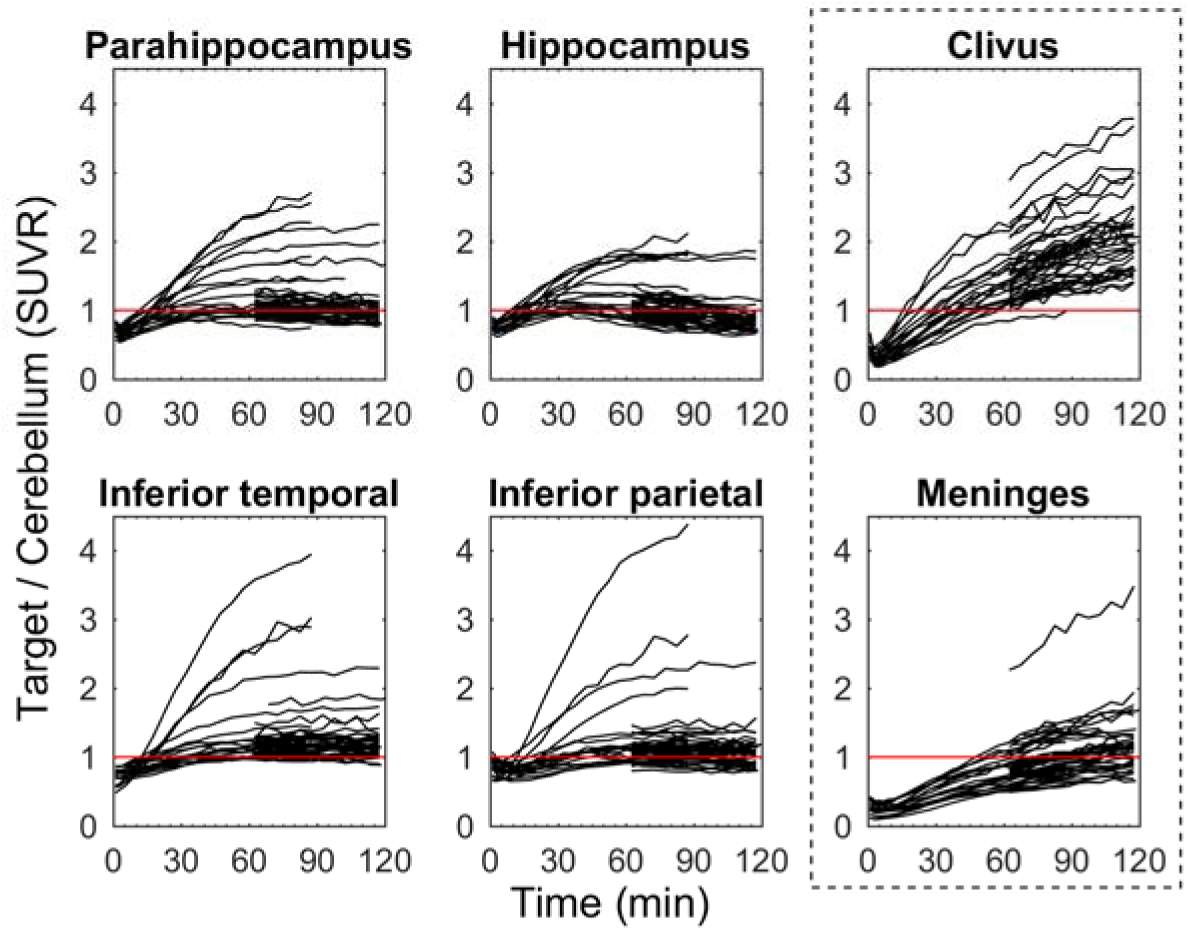
^18^F-MK-6240 Target-to-cerebellum ratio time-activity curves for regions associated with tau pathology (left two columns) and regions with off-target binding (outlined right column) for all 51 participants. The red line indicates an SUVR=1.

### Quantification of ^18^F-MK-6240 Specific Binding

The initial fitting times (t*) for LGA (k_2_=0.04 min^−1^, based on median MRTM2 estimates) and MRTM2 were 35 and 30 minutes, respectively. Regression (table 3) of MRTM2 onto LGA DVR using the full dynamic scan duration was near unity (Fig. 3) with LGA slightly underestimating MRTM2 DVR. Regression outcomes were similar when removing regions with DVR values less than 1.3 from the regression analysis. When shortening the dynamic scan duration, DVR estimates remained stable down to 60 minutes for LGA and 70 minutes for MRTM2 with lower intramethod variability for LGA as compared to MRTM2 for the same scan durations.

**FIGURE 3.**
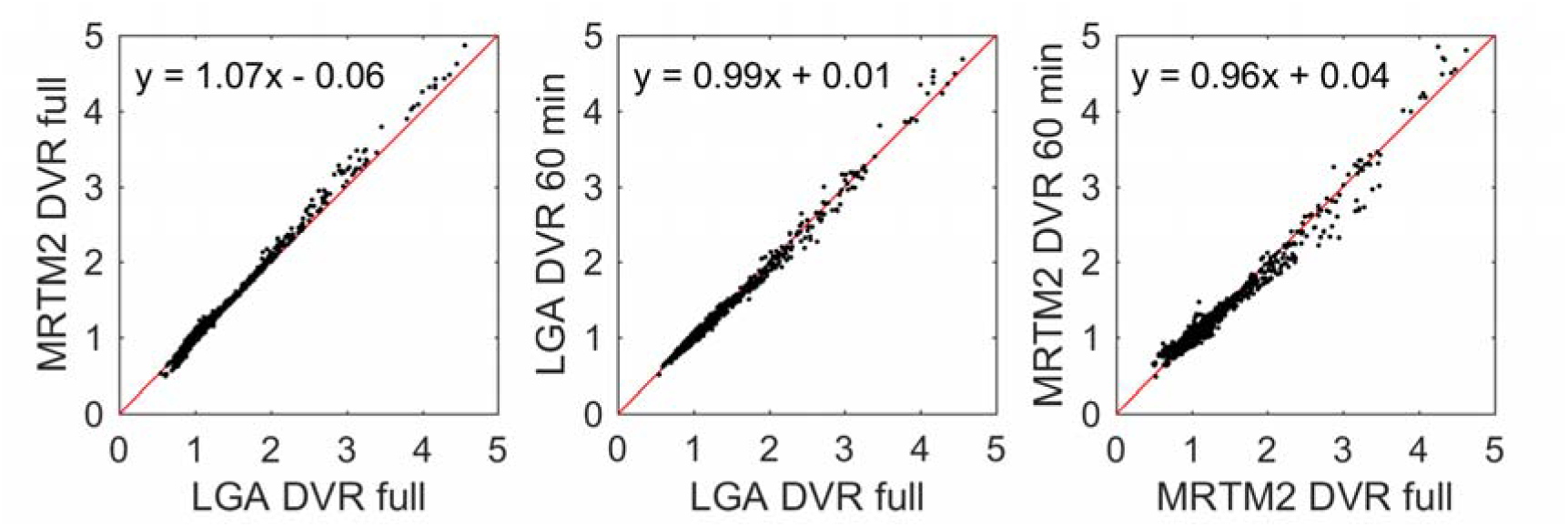
Comparison of ^18^F-MK-6240 DVR estimates using MRTM2 and LGA (left) using the full dynamic time series (90, 105, or 120 minutes), and LGA (middle) and MRTM2 (right) DVR estimates using shortened 60-minute dynamic scans compared to their corresponding full dynamic estimates. Red lines indicate unity (slope = 1, intercept = 0).

**TABLE 3.** Summary of regression statistics for comparisons of DVR methods (top row), DVR using shortened scan durations (rows 2-9), and DVR with stepwise SUVR (rows 10-17). Regressions included all non-cerebellar AAL regions of interest (n=90) and all participants with full dynamic data available (n=19). (SE=standard error, CI=confidence interval, *full* indicates the entire dynamic scan duration (90, 105, or 120 min) was used for DVR estimation)

| Regression | Coefficient | $\beta$ (SE) | 95% CI | R<sup>2</sup> |
| --- | --- | --- | --- | --- |
| *MRTM2 vs. LGA DVR* | Slope | 1.066(0.002) | 1.062,1.069 | 0.995 |
|  | Intercept | -0.060(0.002) | -0.064,-0.055 |  |
| *LGA DVR (50min vs. full)* | Slope | 0.976(0.003) | 0.970,0.983 | 0.981 |
|  | Intercept | 0.019(0.004) | 0.012,0.028 |  |
| *LGA DVR (60min vs. full)* | Slope | 0.986(0.002) | 0.982,0.990 | 0.992 |
|  | Intercept | 0.010(0.003) | 0.005,0.016 |  |
| *LGA DVR (70min vs. full)* | Slope | 0.987(0.001) | 0.984,0.989 | 0.997 |
|  | Intercept | 0.010(0.002) | 0.007,0.013 |  |
| *LGA DVR (80min vs. full)* | Slope | 0.991(0.001) | 0.990,0.992 | 0.999 |
|  | Intercept | 0.008(0.001) | 0.007,0.010 |  |
| *MRTM2 DVR (50min vs. full)* | Slope | 0.841(0.004) | 0.834,0.848 | 0.966 |
|  | Intercept | 0.147(0.005) | 0.138,0.157 |  |
| *MRTM2 DVR (60min vs. full)* | Slope | 0.958(0.004) | 0.951,0.966 | 0.974 |
|  | Intercept | 0.040(0.005) | 0.031,0.050 |  |
| *MRTM2 DVR (70min vs. full)* | Slope | 0.951(0.002) | 0.947,0.955 | 0.993 |
|  | Intercept | 0.051(0.002) | 0.046,0.056 |  |
| *MRTM2 DVR (80min vs. full)* | Slope | 0.964(0.002) | 0.959,0.968 | 0.991 |
|  | Intercept | 0.040(0.003) | 0.034,0.045 |  |
| *SUVR(40-60) vs. LGA(full)* | Slope | 0.814(0.004) | 0.806,0.822 | 0.96 |
|  | Intercept | 0.245(0.005) | 0.235,0.255 |  |
| *SUVR(50-70) vs. LGA(full)* | Slope | 0.934(0.004) | 0.926,0.941 | 0.971 |
|  | Intercept | 0.131(0.005) | 0.122,0.141 |  |
| *SUVR(60-80) vs. LGA(full)* | Slope | 1.014(0.004) | 1.006,1.022 | 0.972 |
|  | Intercept | 0.048(0.005) | 0.038,0.058 |  |
| *SUVR(70-90) vs. LGA(full)* | Slope | 1.073(0.005) | 1.064,1.082 | 0.970 |
|  | Intercept | -0.015(0.006) | -0.026,-0.004 |  |
| *SUVR(40-60) vs. MRTM2(full)* | Slope | 0.757(0.004) | 0.748,0.766 | 0.947 |
|  | Intercept | 0.299(0.006) | 0.288,0.310 |  |
| *SUVR(50-70) vs. MRTM2(full)* | Slope | 0.869(0.004) | 0.861,0.878 | 0.960 |
|  | Intercept | 0.192(0.006) | 0.181,0.203 |  |
| *SUVR(60-80) vs. MRTM2(full)* | Slope | 0.945(0.004) | 0.937,0.954 | 0.964 |
|  | Intercept | 0.112(0.006) | 0.101,0.123 |  |
| *SUVR(70-90) vs. MRTM2(full)* | Slope | 1.002(0.005) | 0.993,1.011 | 0.964 |
|  | Intercept | 0.052(0.006) | 0.040,0.063 |  |

Regression of SUVR onto DVR for 20-minute scans beginning 60 or 70 minutes post-injection (Fig. 4) indicated regression outcomes near unity (Table 3) with SUVR underestimating DVR (LGA and MRTM2) for 20-minute windows starting earlier than 60 minutes. Plots of SUVR onto DVR appeared bi-linear, with values around 1 having a different slope than values above ∼1.5 DVR. Fitting parameters closest to unity between SUVR and MRTM2 were chosen as the criteria for the timing window (70-90 min) used for parametric SUVR image generation.

**FIGURE 4.**
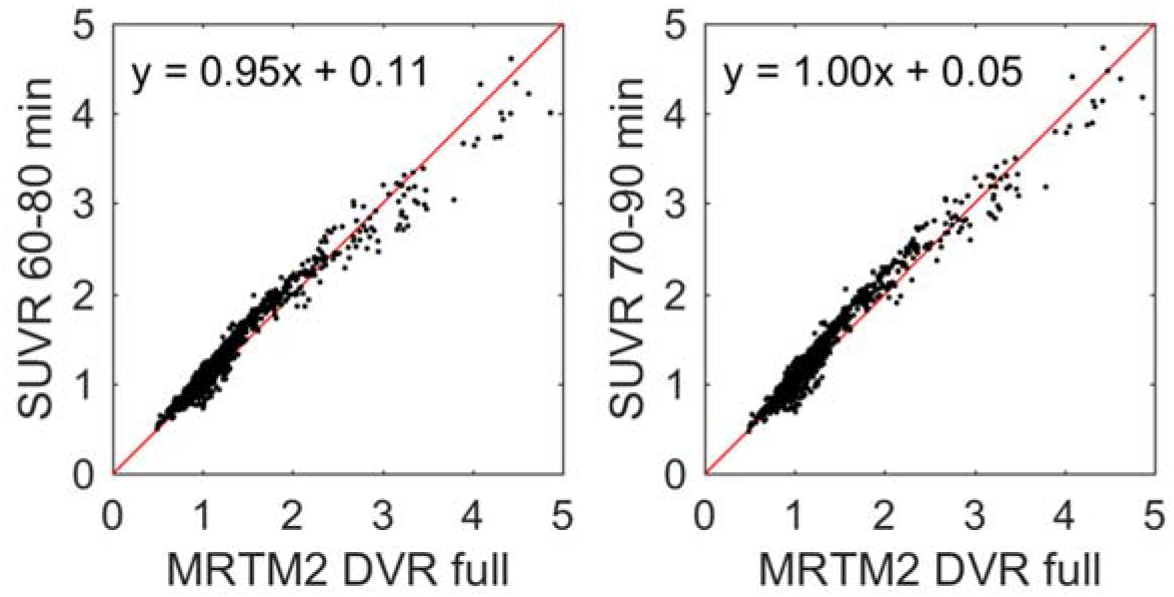
Comparison of SUVR with MRTM2 DVR for SUVR determined from 60-80 (left) and 70-90 (right) minutes post ^18^F-MK-6240 injection. The red line indicates unity.

### ^18^F-MK-6240 Imaging Features

Regions of potential off-target binding identified using mean SUVR images of controls and individual cases included the ethmoid sinus, clivus, sphenotemporal buttress, pineal gland, substantia nigra, superior anterior vermis, superior cerebellum, and the meninges and varied in magnitude and spatial extent across all subjects (Fig. 5). In some extreme cases (6 of 51), meninges binding was observed to spill into adjacent cortical areas. In two cases, focal binding was observed in benign calvarial lesions. Elevated binding was generally not observed in the basal ganglia, choroid plexus (except 1 moderate case), or other regions of the brain that appeared to preclude binding quantification in NFT associated regions. All individuals that were PiB(+) and indicated visually elevated binding in pathological tau-associated regions followed patterns consistent with neuropathological NFT staging (Fig. 6).

**FIGURE 5.**
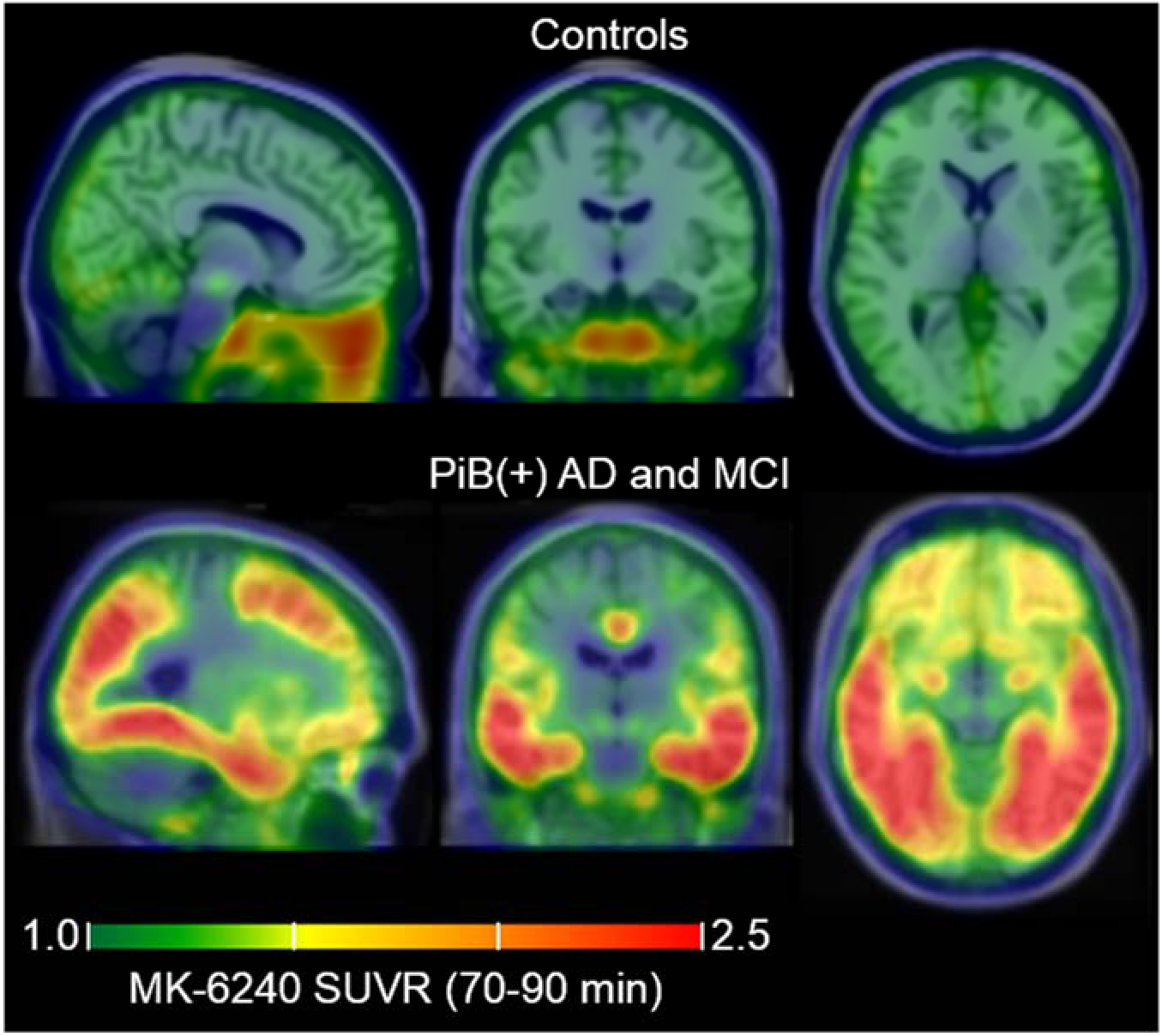
Mean parametric ^18^F-MK-6240 SUVR(70-90 minutes) images taken across controls (top, n=29) and PiB(+) AD and MCI individuals (bottom, n=6) in MNI template space demonstrating common off-target and on-target binding.

**FIGURE 6.**
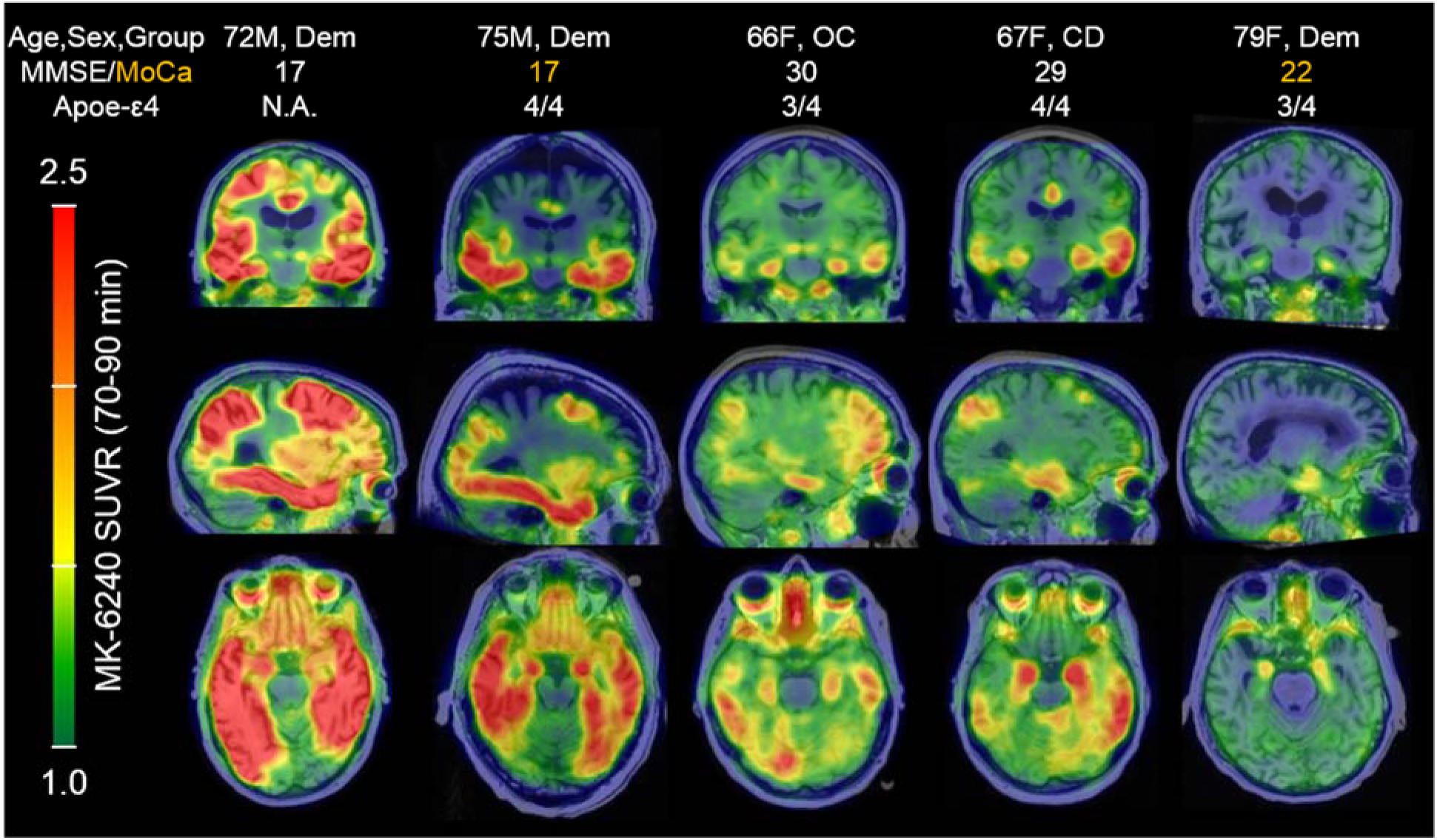
^18^F-MK-6240 parametric SUVR(70-90 min) images in PiB(+) individuals organized by image-based Braak stages. ^18^F-MK-6240 spatial binding patterns in PiB(+) individuals recapitulated patterns consistent with neuropathological staging of Alzheimer’s disease, including in the hippocampus. The PiB(+) dementia case in the far rightward column was clinical diagnosed with probable AD dementia (not informed by biomarkers), but exhibited only circumscribed ^18^F-MK-6240 signal in the entorhinal region.

## DISCUSSION

PET radiopharmaceuticals for detecting NFTs in AD require 1) a high selectivity over other amyloids (e.g. Aβ), 2) high *in vivo* affinity to tau to improve sensitivity for detecting early and longitudinal changes, 3) low off-target binding near regions of interest, and 4) pharmacokinetic properties that enable timely PET acquisition. The kinetics properties of ^18^F-MK-6240 were favorable for PET imaging and comparable to ^18^FAV-1451 (29), a widely used tau PET radiotracer. ^18^F-MK6240 DVR values in AD participants were high (DVR>4) suggesting a combination of high *in vivo* affinity to tau and low non-displaceable signal. Unlike ^18^F-AV-1451 and ^18^F-THK-5351 (12,16), ^18^F-MK-6240 does not appear to have any substantial binding in regions of the brain that would preclude detection of NFTs (e.g. basal ganglia, choroid plexus), particularly in medial temporal regions (e.g. entorhinal cortex and the hippocampus) where tau pathology is implicated relatively early in the disease process (4). In addition, among the subset that were identified as being amyloid positive, ^18^F-MK-6240 binding patterns recapitulated neuropathological staging of NFTs including Braak I and II regions, which supports the sensitivity of the ligand to detect and characterize tau aggregates during early-stage disease. Notably, the lack of off-target binding near the hippocampus (e.g. choroid plexus), a region associated with learning and memory, may allow ^18^F-MK-6240 to differentiate relationships between tau and other features of AD (beta amyloid, atrophy, glucose metabolism, etc.) and their impact on declining cognition.

When evaluating quantification methods for tau PET imaging within the context of clinical research, it is desirable to reduce the PET scan duration to accommodate aging and symptomatic individuals that can experience discomfort, and to maximize the efficiency of the tomograph usage in multi-tracer studies. While DVR estimates were stable using as little as 60 minutes of dynamic data, SUVR quantification with static imaging may be more practical for persons with AD since the overall emission scan duration can be reduced to 20-minutes and still achieve binding estimates comparable to DVR methods.

The selection of the scan duration for SUVR estimation with ^18^F-MK-6240 involves a trade-off between unwanted off-target spill-in from sites outside of the brain with the accuracy of quantification. In particular, regions that could potentially influence cortical binding estimates (ethmoid, clivus and meninges) had SUVRs that were increasing through 120 minutes. Notably, signal in the ethmoid sinus was observed to spill into the orbitofrontal cortex, which could limit quantification of tau-related signal. In contrast, SUVRs in target regions were in agreement with DVR estimates when using data from 70-90 minutes, although SUVRs in target regions were still increasing in higher binding AD subjects during this window. Taken together, this suggests that the 70-90 minute acquisition window will produce accurate binding estimates while reducing potential contamination from off-target binding. This may need to be re-evaluated in studies looking to characterize changes (longitudinal and therapeutic intervention) in SUVR in high binding AD subjects.

A limitation of this study was the absence of arterial blood sampling, which would have provided the gold standard comparison for the DVR and SUVR estimates and could elucidate the source of the discrepancy between LGA and MRTM2 methods. However, since there has been extensive evaluation of the cerebellum as a reference region with other tau tracers (27–30),the cerebellum is used in large-scale tau PET neuroimaging studies (12,13,31), and previous preclinical work did not observe non-polar radiometabolites (18), we believe the results presented in this work will accurately represent comparisons with arterial derived specific binding estimates, but this must be confirmed. Additionally, the mechanism of tracer accumulation in non-NFT target regions is unknown and should be further investigated. Lastly, this study sample was selected to encompass a wide range of disease states, but further investigation of associations between MK-6240 and other disease biomarkers and cognition in a larger cohort is needed.

## CONCLUSION

In a sample of individuals ranging from young cognitively unimpaired controls to individuals with AD dementia, ^18^F-MK-6240 had favorable kinetics for DVR and SUVR quantification by 90-minutes post-injection, low off-target binding in the brain, and binding patterns consistent with neuropathological staging of neurofibrillary tau. These characteristics indicate ^18^F-MK-6240 PET imaging will play a critical role in advancing the understanding of the role of tau pathology in Alzheimer’s disease.

## FUNDING AND ACKNOWLEDGEMENTS

Funding for this work was provided by NIH T32CA009206, NIH R01AG021155 NIH R01AG027161, NIH P50AG033512, and NICHD U54HD090256. The authors would like to acknowledge the University of Wisconsin-Madison Cyclotron Group, the Wisconsin Alzheimer’s Disease Research Center, and the Waisman Center faculty and staff for their contributions to this work and the study participants for their participation and dedication. We would also like to thank Rachel Mulligan of Austin Health for contributing the improved ^18^F-MK-6240 hydrolysis conditions, and Cerveau Technologies for providing the MK-6240 precursor and reference standards.

## DISCLOSURES

[^18^F]MK-6240 precursor and MK-6240 reference standard were provided by Cerveau Technologies. No other potential conflicts of interest relevant to this article exist.

## REFERENCES

1. Braak H, Braak E. Neuropathological stageing of Alzheimer-related changes. Acta Neuropathol. 1991;82:239–259.

2. Thal DR, Rüb U, Orantes M, Braak H. Phases of Aβ-deposition in the human brain and its relevance for the development of AD. Neurology. 2002;58:1791–1800.

3. Nelson PT, Alafuzoff I, Bigio EH, et al. Correlation of Alzheimer disease neuropathologic changes with cognitive status: a review of the literature. J Neuropathol Exp Neurol. 2012;71:362–381.

4. Braak H, Thal DR, Ghebremedhin E, Del Tredici K. Stages of the pathologic process in Alzheimer disease: age categories from 1 to 100 years. J Neuropathol Exp Neurol. 2011;70:960–969.

5. Jack CR, Knopman DS, Jagust WJ, et al. Tracking pathophysiological processes in Alzheimer’s disease: an updated hypothetical model of dynamic biomarkers. Lancet Neurol. 2013;12:207–216.

6. Jack CRJr.,, Wiste HJ, Weigand SD, et al. Defining imaging biomarker cut points for brain aging and Alzheimer’s disease. Alzheimers Dement. 2017;13:205–216.

7. Schwarz AJ, Yu P, Miller BB, et al. Regional profiles of the candidate tau PET ligand 18F-AV-1451 recapitulate key features of Braak histopathological stages. Brain. 2016;139:1539–1550.

8. Grothe MJ, Barthel H, Sepulcre J, et al. In vivo staging of regional amyloid deposition. Neurology. 2017;89:2031–2038.

9. Van Maurik IS, Zwan MD, Tijms BM, et al. Interpreting biomarker results in individual patients with mild cognitive impairment in the Alzheimer’s biomarkers in daily practice (ABIDE) project. JAMA Neurol. 2017;74:1481–1491.

10. Mathis CA, Lopresti BJ, Ikonomovic MD, Klunk WE. Small-molecule PET tracers for imaging proteinopathies. Semin Nucl Med. 2017;47:553–575.

11. Brier MR, Gordon B, Friedrichsen K, et al. Tau and Aβ imaging, CSF measures, and cognition in Alzheimer’s disease. Sci Transl Med. 2016;8:338ra366.

12. Johnson KA, Schultz A, Betensky RA, et al. Tau positron emission tomographic imaging in aging and early Alzheimer disease. Ann Neurol. 2015;79:110–119.

13. Lowe VJ, Wiste HJ, Senjem ML, et al. Widespread brain tau and its association with ageing, Braak stage and Alzheimer’s dementia. Brain. 2018;141:271–287.

14. Villemagne VL, Dore V, Bourgeat P, et al. Abeta-amyloid and tau imaging in dementia. Semin Nucl Med. 2017;47:75–88.

15. Shoghi-Jadid K, Small GW, Agdeppa ED, et al. Localization of neurofibrillary tangles and beta-amyloid plaques in the brains of living patients with Alzheimer disease. Am J Geriatr Psychiatry. 2002;10:24–35.

16. Ng KP, Pascoal TA, Mathotaarachchi S, et al. Monoamine oxidase B inhibitor, selegiline, reduces ^18^F-THK5351 uptake in the human brain. Alzheimers Res Ther. 2017;9:25.

17. Hashimoto H, Kawamura K, Takei M, et al. Identification of a major radiometabolite of [11C]PBB3. Nucl Med Biol. 2015;42:905–910.

18. Hostetler ED, Walji AM, Zeng Z, et al. Preclinical characterization of ^18^F-MK-6240, a promising PET tracer for in vivo quantification of human neurofibrillary tangles. J Nucl Med. 2016;57:1599–1606.

19. Johnson SC, Koscik RL, Jonaitis EM, et al. The Wisconsin Registry for Alzheimer’s Prevention: A review of findings and current directions. Alzheimers Dement(Amst). 2018;10:130–142.

20. Johnson SC, Christian BT, Okonkwo OC, et al. Amyloid burden and neural function in people at risk for Alzheimer’s Disease. Neurobiol Aging. 2014;35:576–584.

21. Lazari M, Quinn KM, Claggett SB, et al. ELIXYS - a fully automated, three-reactor high-pressure radiosynthesizer for development and routine production of diverse PET tracers. EJNMMI Res. 2013;3:52.

22. Betthauser TJ, Ellison PA, Murali D, et al. Characterization of the radiosynthesis and purification of [18F]THK-5351, a PET ligand for neurofibrillary tau. Appl Radiat Isot. 2017;130:230–237.

23. Racine AM, Clark LR, Berman SE, et al. Associations between performance on an abbreviated CogState battery, other measures of cognitive function, and biomarkers in people at risk for Alzheimer’s disease. J Alzheimers Dis. 2016;54:1395–1408.

24. Christian BT, Vandehey NT, Floberg JM, Mistretta CA. Dynamic PET denoising with HYPR processing. J Nucl Med. 2010;51:1147–1154.

25. Logan J, Fowler JS, Volkow ND, Wang G-J, Ding Y-S, Alexoff DL. Distribution volume ratios without blood sampling from graphical analysis of PET data. J Cereb Blood Flow Metab. 1996;16:834–840.

26. Ichise M, Liow JS, Lu JQ, et al. Linearized reference tissue parametric imaging methods: application to [^11^C]DASB positron emission tomography studies of the serotonin transporter in human brain. J Cereb Blood Flow Metab. 2003;23:1096–1112.

27. Lemoine L, Saint-Aubert L, Marutle A, et al. Visualization of regional tau deposits using ^3^HTHK5117 in Alzheimer brain tissue. Acta Neuropathol Commun. 2015;3:40.

28. Marquié M, Normandin MD, Vanderburg CR, et al. Validating novel tau positron emission tomography tracer [F-18]-AV-1451 (T807) on postmortem brain tissue. Ann Neurol. 2015;78:787–800.

29. Wooten DW, Guehl NJ, Verwer EE, et al. Pharmacokinetic evaluation of the tau PET radiotracer ^18^F-T807 (^18^F-AV-1451) in human subjects. J Nucl Med. 2017;58:484–491.

30. Jonasson M, Wall A, Chiotis K, et al. Tracer kinetic analysis of (S)-^18^F-THK5117 as a PET tracer for assessing tau pathology. J Nucl Med. 2016;57:574–581.

31. Gordon BA, Friedrichsen K, Brier M, et al. The relationship between cerebrospinal fluid markers of Alzheimer pathology and positron emission tomography tau imaging. Brain. 2016;139:2249–2260.

